# Urban Environments Reshape Reproductive Phenology in Plants Across the Tropics

**DOI:** 10.64898/2026.01.28.702306

**Authors:** Rohit Raj Jha, Anita Simha, Richard Ekeng Ita, Rachana Rao, Daijiang Li, Gaurav Kandlikar

## Abstract

Plant phenological responses to global change phenomena like urbanization remain understudied in the tropics, hindering predictions regarding the dynamics of tropical ecosystems amid rapid land use changes. Studies of tropical phenology are limited by complexities, like the limited availability of phenological data, especially in urbanized landscapes. Observations recorded on citizen science platforms can overcome this limitation by providing vast, spatially distributed data. In this study, we utilize iNaturalist data to evaluate plant reproductive phenology in tropical urban vs. rural habitats. We first compare iNaturalist data (111533 records) to herbarium collections (217991 records) in order to validate their use, and we then investigate urban-rural phenology differences within 25-km spatial grids for 238 species. Data from iNaturalist and herbaria yield complementary insights, with the former being uniformly distributed between urban and rural settings, and the latter biased towards rural observations. On average, we found species to have significantly longer reproductive duration (β = 11.79 ± 2.83 SE, t = 4.16, p < 10^4), and correspondingly weaker strength of seasonality in urban settings than in nearby rural localities. We also find trait-mediated variation, with seasonal, annual, and herbaceous plants showing more pronounced differences in reproductive duration and seasonality strength. These results suggest that urbanization in tropical landscapes might have important implications for plant demography, with potential consequences for community and ecosystem dynamics. Our work also points to the value of integrating insights from natural history collections with data from citizen science platforms for enabling broad-scale insights into ecological dynamics in tropical urban landscapes.

## 1. INTRODUCTION

Phenology, the study of the timing of recurring biological events, links organisms to their abiotic environment, including temperature, precipitation, and photoperiod, and underpins ecosystem functioning (Caparros-Santiago et al., 2021; Johansson et al., 2015). In terrestrial systems, the timing of events such as leaf-out, flowering, and fruiting in plants can influence primary productivity, nutrient cycling, and energy flow within ecosystems (Forrest & Miller-Rushing, 2010; Gallinat et al., 2021; Moore et al., 2016; Tang et al., 2016), as well as ecological interactions like pollination, herbivory, predation, and competition (Forrest & Miller-Rushing, 2010; Johansson et al., 2015; Kharouba et al., 2018). Shifts in plant phenology are among the earliest and most consistent biological responses to global environmental change, serving as key indicators of species’ adaptive capacity and ecosystem resilience (Cleland et al., 2007; Parmesan & Yohe, 2003). Most current understanding of these impacts comes from temperate regions, where seasonal variations in important cues like climate and daylength are well-defined across the year. For example, Menzel et al., (2006)’s landmark study linking phenology to climate warming relied on phenological data from a systematic network across Europe to show that a vast majority of plant species show signals of advanced leafout, flowering, and fruiting, consistent with warmer winter temperatures and earlier spring conditions. Climate-driven shifts in plant phenology have now been identified across temperate regions (Stuble et al., 2021), although the dynamics and drivers of these shifts can vary across continents (Zohner et al., 2017). Conversely, much less is known about phenological responses to global environmental change in tropical ecosystems (Piao et al., 2019), where rainfall patterns and uniform temperatures give rise to complex, multi-modal seasonal patterns (Abernethy et al., 2018; Fitchett et al., 2015).

Among the most prominent forces for environmental change in the tropics is urbanization, which is associated with changes to local temperature patterns, water availability, and biological diversity (Kabano et al., 2021; McDonald et al., 2020; Ogunbode et al., 2025; Ribeiro et al., 2024). Studies of urban phenological shifts are useful for identifying urban heat islands and evaluating impacts of warming on phenology (Jochner et al., 2013). In temperate cities, urban heat islands and irrigation systems often reduce climatic constraints on plant development, leading to earlier or extended reproductive periods in cities (Li et al., 2019; Neil & Wu, 2006; D. S. Park et al., 2023; Wohlfahrt et al., 2019). Although urbanization has been rapid and widespread throughout the tropics and is of particular concern for tropical biodiversity conservation (McDonald et al., 2020; Simkin et al., 2022), tropical cities are surprisingly understudied in phenological research (Bonebrake et al., 2025), which constrains predictions regarding the resilience of tropical plant dynamics amid rapid climatic and land use changes. Understanding how urbanization influences reproductive phenology in the tropics is therefore a pressing research gap (Kabano et al., 2021; Marcacci et al., 2023).

Several challenges complicate studies of tropical phenology. First, unlike in temperate regions, where strong seasonal temperature and photoperiod cues trigger synchronized phenological events, tropical phenology is influenced by a broader suite of environmental cues such as rainfall, irradiance, and local microclimate (Borchert, 1996; Borchert et al., 2002; Jochner et al., 2013; Numata et al., 2022). These cues may operate asynchronously, and their timing, strength, and biological relevance can vary across space, years, and species phenology (Abernethy et al., 2018; J. Y. Park et al., 2019; Sakai, 2001; Singh & Kushwaha, 2005). Second, tropical ecosystems are generally more species-rich than temperate regions, which results in tropical plant communities encompassing a wider variety of phenological strategies (Abernethy et al., 2018; J. Y. Park et al., 2019; Singh & Kushwaha, 2005). Along with this, plants’ life history and growth form also play a definite role in phenological responses, as annual and perennial species and contrasting growth forms vary in resource allocation strategies and dependency on environmental cues (Borchert, 1996; Sakai, 2001). Finally, the study of tropical phenology has historically been constrained by the limited availability of long-term phenological monitoring plots and field stations (Abernethy et al., 2018; Bush et al., 2017, 2018). Although remote sensing techniques can capture seasonal changes in canopy greenness, they fall short in capturing species- and population-level phenology (Fisher et al., 2006; Zhang et al., 2006) (Fisher et al., 2006; Zhang et al., 2006).

One potential approach to overcome these limitations is through using data derived from citizen science platforms, which have the potential to transform phenology research by providing vast, globally distributed datasets of species occurrences. These data are timestamped, georeferenced, and often include photographs that enable the inference of phenological events across unprecedented spatial and temporal scales (Barve et al., 2020; Callaghan et al., 2020; Wolf et al., 2022). For instance, iNaturalist, a widely used citizen science platform, currently hosts over 69 million research grade plant observations. 4.6 million of these observations come from tropical Asia, a traditionally data-scarce region, and this number is increasing rapidly (Di Cecco et al., 2021). In temperate ecosystems, iNaturalist records have been proven to yield valuable insights into plant phenology. For example, Li et al. (2019) used 22 million images to extract phenology records from the USA and Europe and found that urbanization advances flowering and leaf-out in colder regions. Similarly, (Iwanycki Ahlstrand et al., 2022) compared directed citizen science, herbarium, and iNaturalist records for three spring flowering species in Denmark, and found that iNaturalist provided the broadest spatial coverage and captured peak flowering well. Moreover, combining citizen science dataset with traditional monitoring networks and herbarium records can help fill spatial and temporal gaps, enhancing the detection of climate-driven phenological changes (Davis et al., 2015; D. S. Park et al., 2023; Willis et al., 2017). In sum, despite important limitations such as observer bias, variable sampling effort, and uneven taxonomic coverage, opportunistically collected observation records in citizen science platforms can yield meaningful phenological insights, especially when analyzed with modern circular statistics and hierarchical modeling techniques (Capinha et al., 2024; Lai, 2025; Pabon-Moreno et al., 2019; Willig et al., 2024).

In this study, we utilize iNaturalist data to evaluate plant reproductive phenology in urban vs. rural habitats across the tropical latitudes. We first evaluated the value of iNaturalist observations for tropical phenological research by comparing phenological estimates derived from these observations to those derived from herbarium specimens collected in tropical latitudes. Next, we asked whether plant reproductive phenology in urban centers differs from reproductive phenology in adjacent rural areas. Finally, we evaluated whether plant growth and reproductive strategies impact phenological responses to urbanization. Our study offers one of the first large-scale, trait-specific evaluations of how urbanization influences reproductive phenology in tropical plants.

## 2. METHODS

### 2.1. Data acquisition

#### 2.1.1 Plant observations from iNaturalist

As a source of our citizen science data, we used plant data collected through iNaturalist (Fig. 1, left panel). To retrieve observations along with their phenology status (presence of flowers, fruits, or both), we accessed data through PhenoBase (Dinnage et al., 2025). PhenoBase, through its machine learning system known as PhenoVision, has processed millions of field photos from iNaturalist and provides them with phenology status and geographic and temporal metadata. PhenoVison detects flowers and fruits using a Vision Transformer (ViT) model fine-tuned with a masked autoencoder pretraining method. This machine learning program was trained on over 1.5 million human-annotated iNaturalist images, achieving high validation accuracy (98.5% for flowers, 95% for fruits). Additional calibration steps refined detection thresholds and reduced false positives, ensuring high-confidence phenological observations. As annotations on PhenoBase are limited to observations between 2010 and 2023, we limited our search to those years. We downloaded 217,991 observations representing 296 species that had a minimum of 100 observations within the tropics (23.5^0^ N – 23.5^0^ S) during the defined time period.

**Figure 1.**
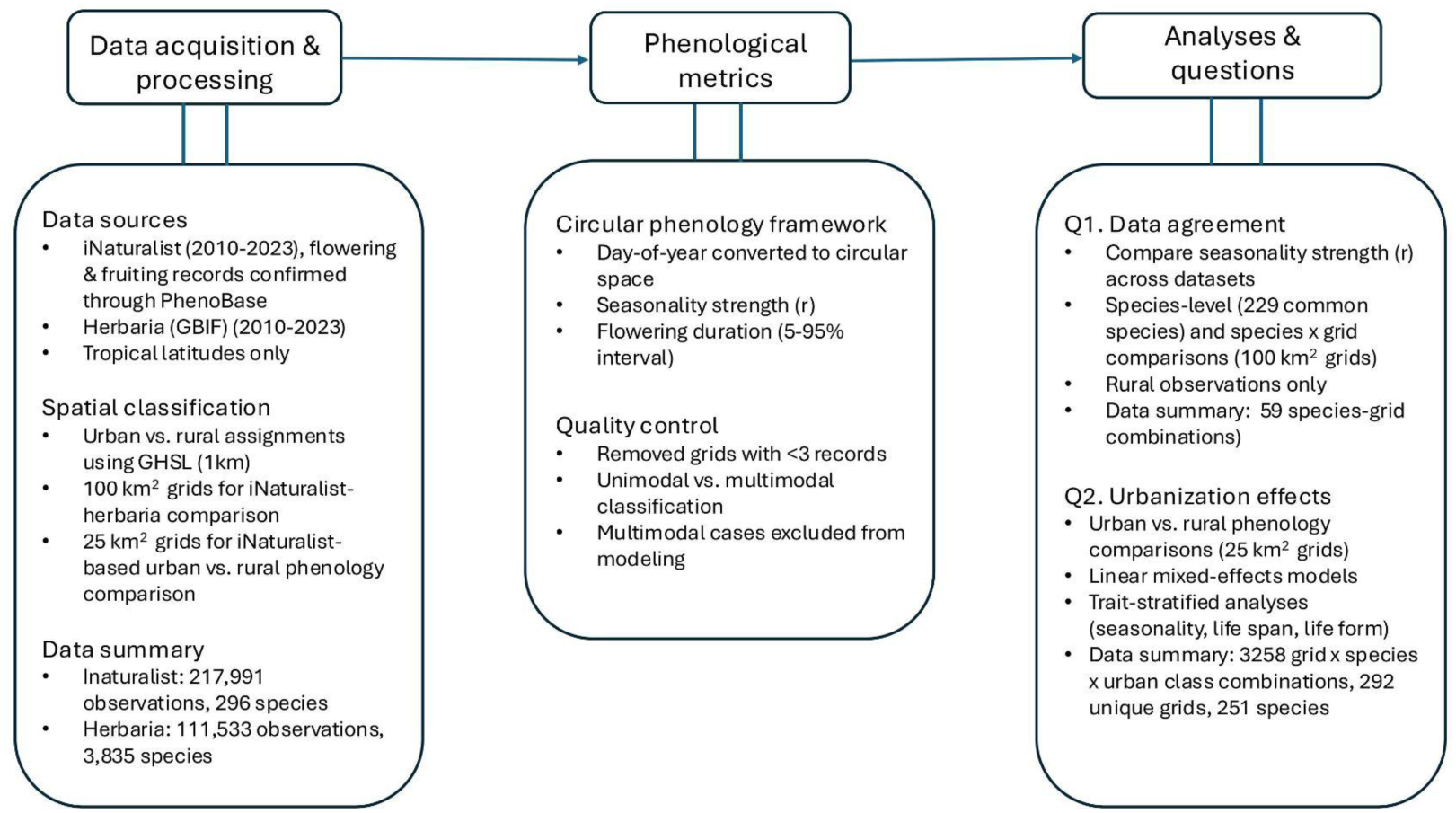
Overview of data source, phenological metrics, and analytical framework used to assess urban-rural differences in tropical plant reproductive phenology

#### 2.1.2. Plant observations from herbarium records

We retrieved data from the Global Biodiversity Information Facility (GBIF) to compare phenological estimates derived from observations on iNaturalist to those derived from tropical plant specimens deposited into herbaria worldwide (Fig. 1, left panel). We queried GBIF for all plant specimen records from tropical latitudes collected between 2010-2023, with the additional constraints that the record was not marked as having a geospatial issue and had collection coordinates with uncertainty of less than 100m. To expand the geographic and taxonomic scope of this herbarium-derived data, we also retrieved tropical herbarium records from Tropicos, a curated botanical database maintained by the Missouri Botanical Garden; this dataset had not been retrieved with our previous query as it has unspecified coordinate uncertainties. Together, these queries yielded 111533 observations from 3853 species with a minimum of 10 records collected globally within 2010-2023. Subsequent analyses assume that the presence of a herbarium record indicates that the plant in question was reproductive (i.e., flowering or fruiting) at the time of collection, based on common collection practices that promote collecting plants with reproductive features present (Goëau et al., 2020; Heberling et al., 2019; Willis et al., 2017).

#### 2.1.3 Urbanization

To assign urbanization status for each observation, we used the 2020 version of the Global Human Settlement-Degree of Urbanization (GHS-SMOD) layer, developed by the European Commission’s Joint Research Centre (Fig. 1, left panel) (Florczyk et al., 2019). This layer offers consistent global data on human settlement intensity at a 1 km resolution, based on population density and built-up areas. Each grid cell is classified into one of eight urbanization categories, ranging from dense city centers to largely uninhabited, very low-density zones, allowing a continuous representation of settlement intensity. We simplified these categories into two categories: urban (Grid Codes 21, 22, 23, 30: representing peri-urban to urban cores) and rural (Grid Codes 11–13: covering very low-density rural to rural clusters), and assigned an urbanization class to each plant observation (Melchiorri et al., 2018, 2019; Santillan & Heipke, 2023).

### 2.2. Data analysis

#### 2.2.1 Comparison between phenology estimation

To determine if species-level reproductive seasonality from citizen-science data aligns with herbarium-based inferences, we compared observations from iNaturalist and herbaria. For each species in each dataset (iNaturalist and herbarium), day-of-year (DOY) values were converted to radians and summarized using circular statistics in the circular package in R (Lund et al., 2025). We estimated the mean vector length (*r*), a concentration parameter from 0 to 1 that measures the strength of seasonality (Fig. 1, middle panel) (Morellato et al., 2009) and used this to classify the strength of seasonality for each species (Weak (*r* < 0.35), Moderate (0.35 ≤ *r* < 0.70), and Strong (*r* ≥ 0.70)). We selected this *r*-based classification because *r* directly measures effect size, indicating how closely observations cluster around the mean flowering date, regardless of sample size (Alsammani et al., 2023; Staggemeier et al., 2020; Willig et al., 2024). Moreover, when comparing the same species across two large, uneven datasets (GBIF vs. iNaturalist), p-values from circular significance tests can be misleading, as the Rayleigh p-value is highly sample-size-dependent (Alsammani et al., 2023; Willig et al., 2024). We then used a confusion matrix to compare the agreement in flowering strength between two datasets.

#### 2.2.2 Using iNaturalist observations to compare rural vs. urban phenology

##### Exclusion of multimodal species

In the tropics, some species can express multimodal reproductive phenology, with multiple flowering peaks per year (Wright et al., 2019), which complicate comparisons of phenology across urban and rural contexts. To account for this, we first used the Hermans-Rasson (HR) test, a robust, nonparametric method for detecting nonuniformity that does not depend on a specific distribution (Hermans & Rasson, 1985; Landler et al., 2019) and excluded species that express multimodal reproductive phenology from subsequent analyses. First, we used Rayleigh’s test to determine whether observations are significantly clustered around a mean direction (Morellato et al., 2009). Next, we used the Rayleigh test to evaluate the strength of seasonality (aseasonal, when Rayleigh p > 0.05, or seasonal, Rayleigh p < 0.05). Both seasonal and aseasonal species were included in the analysis because the seasonality-strength measure r is relevant for unimodal distributions and interpretable as “weak seasonality” in uniform cases. However, species with multimodal reproductive phenology were excluded, as these violate the assumptions behind the circular statistics used in our analysis (Datta et al., 2025; Morellato et al., 2009; Staggemeier et al., 2020).

##### Modeling reproductive seasonality strength and duration

Reproductive seasonality strength (*r*) was estimated from circular statistics (Fig. 1, right panel). We estimated total reproductive duration by calculating the percentiles along the circular ordering of flowering and fruiting angles and taking the angular distance between the 5th and 95th percentiles, and finally converting this angular distance back into days of year.

Next, to test whether seasonality and duration differ between urban and rural environments, we fitted the following linear mixed-effects models:

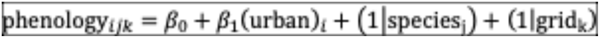

Here, phenology ijk was either a species i’s estimated $r$ or its estimated reproductive duration in grid cell *j* and in urban class *k*. β₁ captures the fixed effect of urbanization as rural and urban class, and species and grid were treated as random intercepts to account for taxonomic and spatial non-independence.

Finally, as the circular statistics in our modeling approach can be sensitive to the number of observations being used to generate the statistic (e.g. longer flowering durations or reduced seasonality strength could be estimated simply due to more extensive sampling), we additionally conducted a supplemental analysis in which we subsampled the data to have equal representation within each species-grid-urban class category (Fig. S5).. Briefly, we “rarefied” the data 1000 times to ensure equal representation in each urban class within a species-grid cell combination, estimated phenology using the circular statistics above, and fitted the global linear mixed effect model. As the results from this approach were consistent with the results of our “global” model that included all observations (Fig. S5), we focus our main text on results from the global model.

##### Modeling with species’ traits

To assess how different plant growth and reproductive strategies impact phenological responses to urbanization, we also modeled: (1) seasonality class, including seasonal and aseasonal species, (2) life duration categories, such as annual and perennial species, and (3) life forms, namely herbs, shrubs, and vines separately for smaller subsets of the data. Trait-specific information for all the species was compiled through manual online searches. We only consulted the authoritative botanical sources like the USDA and botanical gardens’ websites. We fitted models with interactive effects of urbanization class and plant traits (seasonality, life duration, or life forms). All the models were fitted using lmerTest with Satterthwaite-adjusted t-tests, weighting observations by sample size per grid (Kuznetsova et al., 2017). All analyses were performed using R version 4.5.1 (R Core Team, 2025).

## 3. RESULTS

### 3.1 Data overview

We had 217,991 unique observations with flowers or fruits or both from iNaturalist and 111,533 from herbaria, spanning 2010-2023. Our iNaturalist and herbaria datasets had 296 and 3,835 unique species, respectively. They had 229 species in common.

Our iNaturalist dataset (Fig. 2, top panel) showed broad tropical coverage across continents, with dense clusters of observations in South and Southeast Asia, East Africa, and Central America, with roughly equal distributions between rural and urban regions (Fig 2a). In contrast, the herbarium data obtained from GBIF (Fig. 2, bottom panel) were spatially sparser, concentrated mainly in Latin America, Northern Australia, West Africa, and parts of Southeast Asia. Very few herbarium records (∼3.3%) were collected in urban locations (Fig. 2b), consistent with the traditional botanical survey bias toward less disturbed ecosystems. However, herbarium records spanned 3,835 species in contrast to the 229 species represented in iNaturalist, indicating that iNaturalist offers broader spatial coverage with comparable number of observations in urban and rural areas, while herbaria provide taxonomically rich, rural-biased observations.

**Figure 2.**
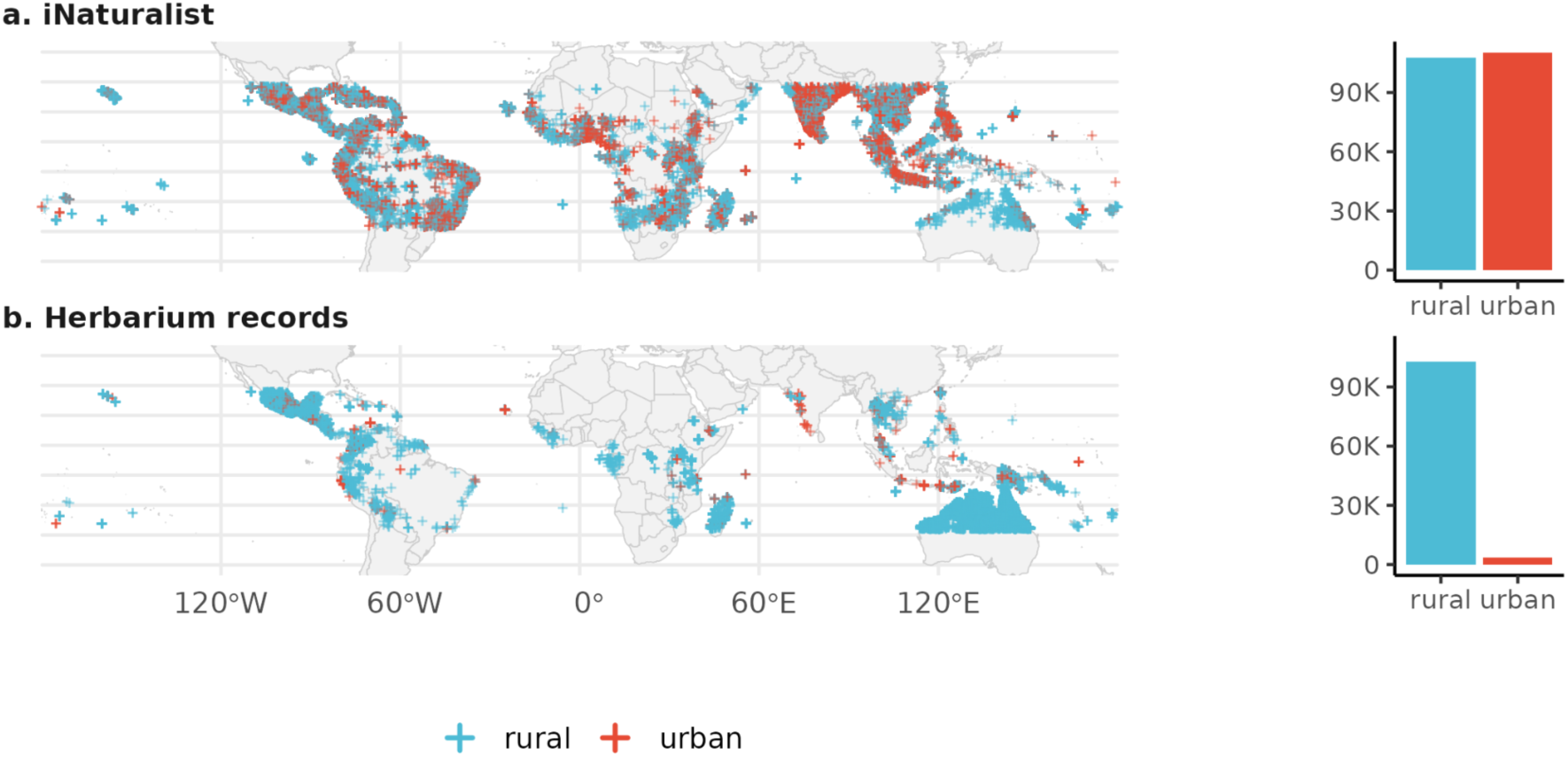
Spatial distribution of observations included from iNaturalist (top row) and herbaria (bottom row) across tropical regions. Colored points represent urban (red) and rural (blue) observations. iNaturalist data exhibit broad spatial coverage with strong urban clustering, whereas herbaria data show a rural-dominated sampling pattern. Inset bars represent total counts of rural and urban observations and are scaled consistently across both figure panels.

**Figure 3.**
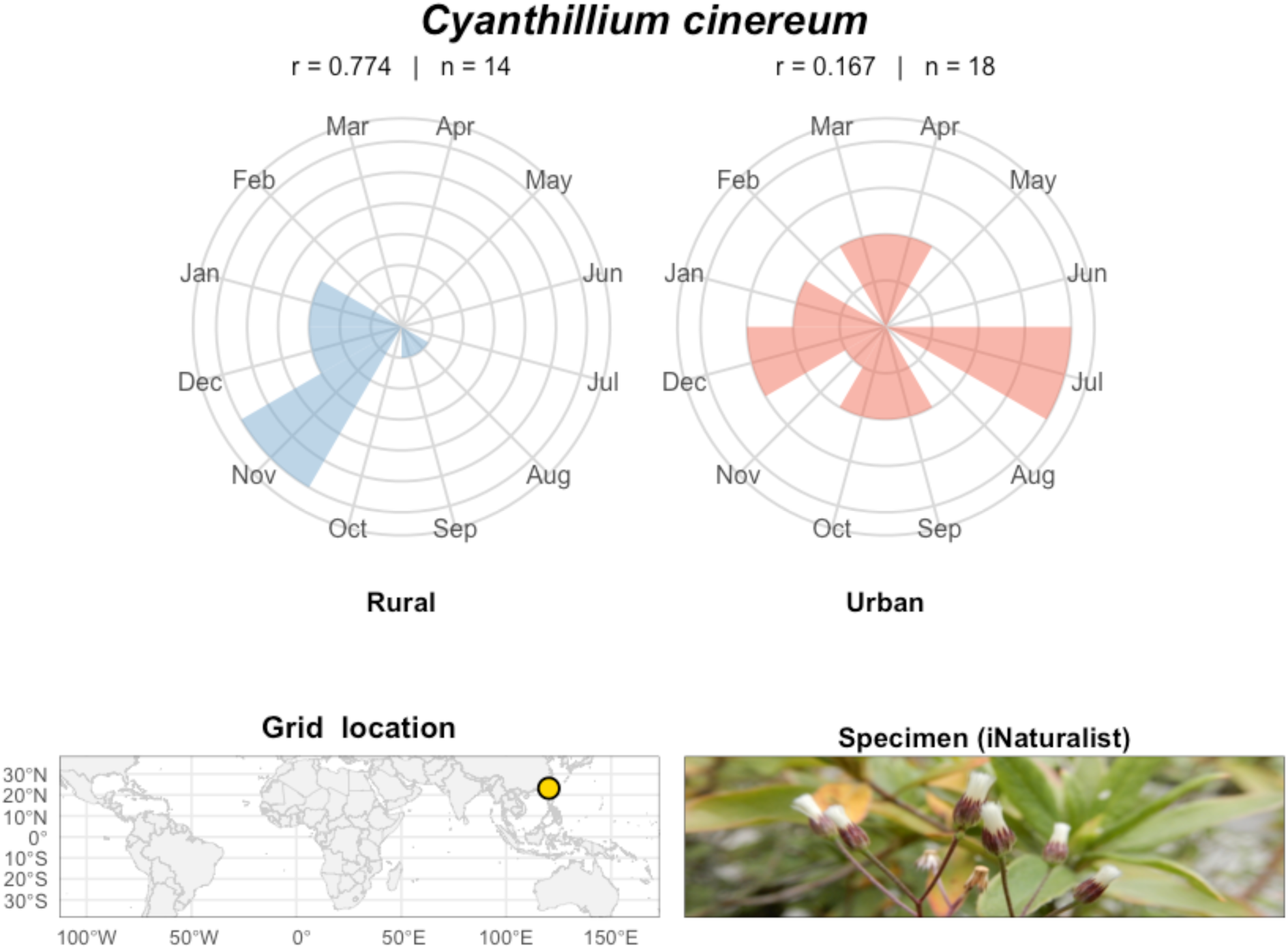
Circular flowering phenology for *Cyanthillium cinereum* (Bottom right) developed using iNaturalist observations from 2010 to 2023, covering a single grid (Bottom left). Rural iNaturalist records show a more concentrated, seasonal flowering peak, whereas urban observations display a broader, less seasonal distribution. r indicates the corresponding values for reproductive seasonality strength, and n is the number of observations in each urban class. Specimen credit: Lava Chen, 2026

Similarly, we found iNaturalist observations peaking in April–May (late dry to early wet season in many tropical regions), reflecting heightened flowering visibility and user activity. Similarly, we observed secondary peaks in October–November, suggesting a bimodal phenological rhythm consistent with intertropical seasonal transitions. Both urban and rural classes show similar seasonal timing, but urban peaks were slightly higher in April, May, and November, reflecting more observer activity and maybe prolonged reproductive activity in cities due to heat islands (Supplementary file, Fig. 1, top panel). Whereas the herbaria dataset with a consistent rural-dominated signal showed a peak collection timing between March and June, and moderate activity throughout the year. The absence of strong urban representation mainly reflects sampling bias rather than biological inactivity (Supplementary file, Fig. 1, bottom panel).

### 3.2 Phenological insights from iNaturalist and herbarium-derived

Classifications of reproductive seasonality were broadly consistent across the herbarium-derived and iNaturalist datasets. Of the 229 species in this comparison, 118 were identified as weakly seasonal with both iNaturalist and herbarium data, and only 7 were identified as strongly seasonal with both datasets (Table 1, top panel). The most common discrepancy occurred due to species being assigned as having “moderate” seasonality through herbarium data and only “weak” seasonality when phenology was assessed with iNaturalist records. Values of the seasonality metric (*r*) from herbarium and iNaturalist records were significantly correlated with one another (Spearman’s ρ = 0.31, p = 1.31 × 10⁻⁵).

**Table 1.** Confusion matrix comparing reproductive seasonality classifications based on data from iNaturalist (rows) and herbaria (columns). The top half shows species-level comparisons, including only taxa present in both datasets. The bottom half shows grid-level data based on shared species × grid combinations across iNaturalist and herbaria. The diagonal cells’ value, denoted in bold shows agreement between datasets.

| iNat / Herbaria | Weak | Moderate | Strong |
| --- | --- | --- | --- |
| Species-level seasonality classification |  |  |  |
| Weak | **118** | 53 | 3 |
| Moderate | 17 | **24** | 2 |
| Strong | 0 | 3 | **6** |

### 3.3 Comparison of reproductive phase duration and reproductive seasonality strength

#### 3.3.1. Global comparison

Across 3104 grid × species × urban class combinations, representing 285 unique grids and 238 species, reproductive duration ranged from 3 to 364 days. On average across species and grid cells, plant reproductive periods were significantly longer in urban than in rural contexts (Fig. 4A; rural flowering period = 195.04 ± 5.06; urban flowering period = 206.83 ± 5.06; p = 3.26e-05, Table S1). The strength of seasonality (*r*) was correspondingly lower in urban than in rural settings (Fig. 4B; rural r = 0.61 ± 0.014; urban r = 0.59 ± 0.014; p = 3.1e-04, Table S2).

**Figure 4.**
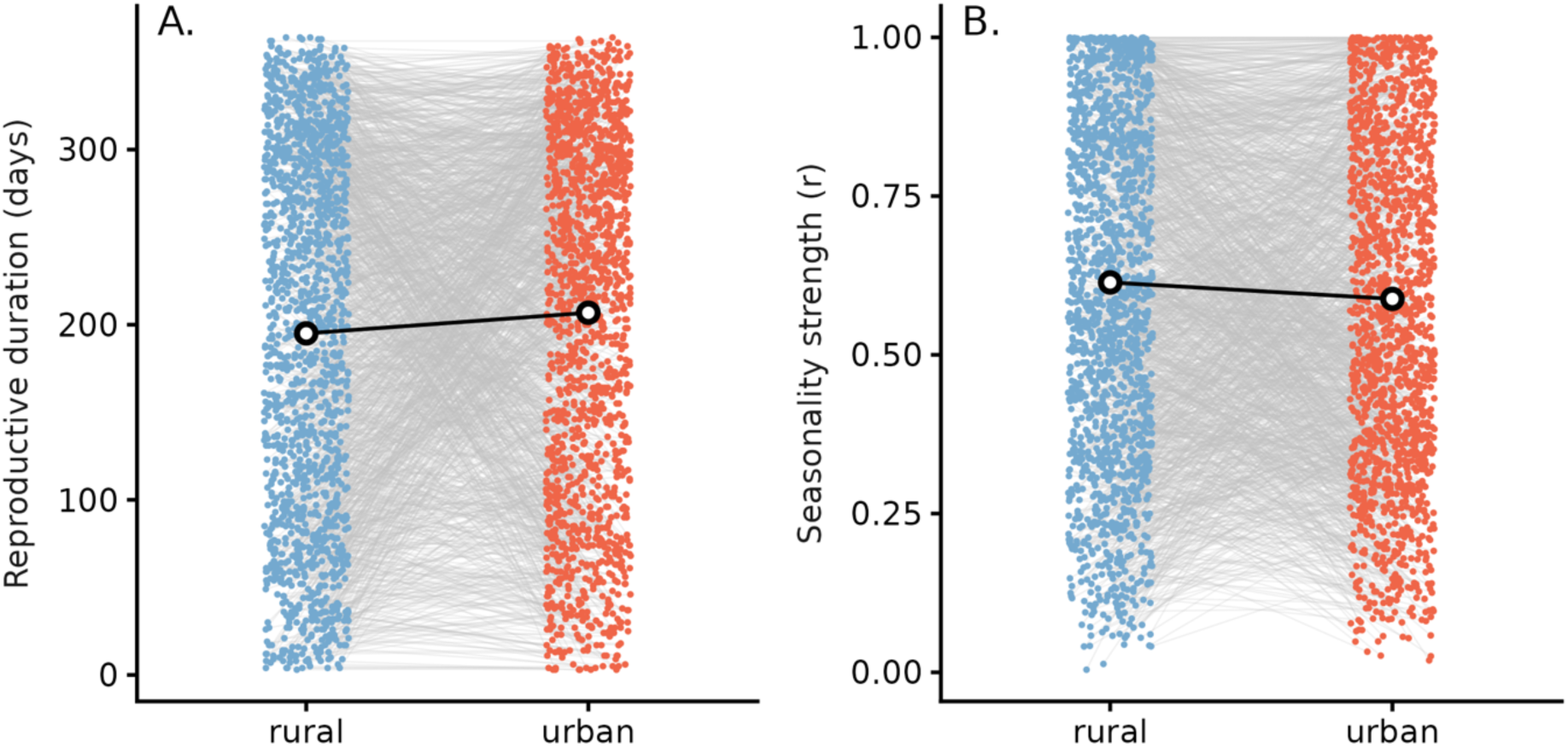
Urban–rural differences in reproductive duration (A) and seasonality strength (B) across all species x grids x urban class combinations. Each colored point represents one species in a grid cell, in either rural (blue) or urban (red) localities; thin grey lines connect urban and rural estimates. The white points and lines represent model-estimated marginal means for rural and urban localities. This result of longer reproductive periods in urban than in rural conditions was supported in a supplemental analysis that controlled for potential effects of uneven urban vs. rural sampling intensities within grids.

#### 3.3.2. Comparison across seasonal and aseasonal species

For species with seasonal reproduction (i.e., those with Raleigh p < 0.05, Datta et al., 2025), reproductive periods were three weeks longer in urban than in rural settings (rural reproductive duration = 177.57 days ± 5.56; urban reproductive duration = 198.78 ± 5.35; p = 5.12 e-15; Fig. 5A and Table S1). Species with aseasonal reproduction had on average longer reproductive periods; these species also experienced extended reproductive periods in urban than rural settings, although this effect was weaker than for seasonal species (rural reproductive duration = 213.65 days ± 5.42; urban reproductive duration = 222.57 ± 5.35; p = .03; Fig. 5B and Table S1). Similar results were also detected when measuring seasonality based on *r* rather than flowering duration (Table S2 and Fig. S2).

**Figure 5.**
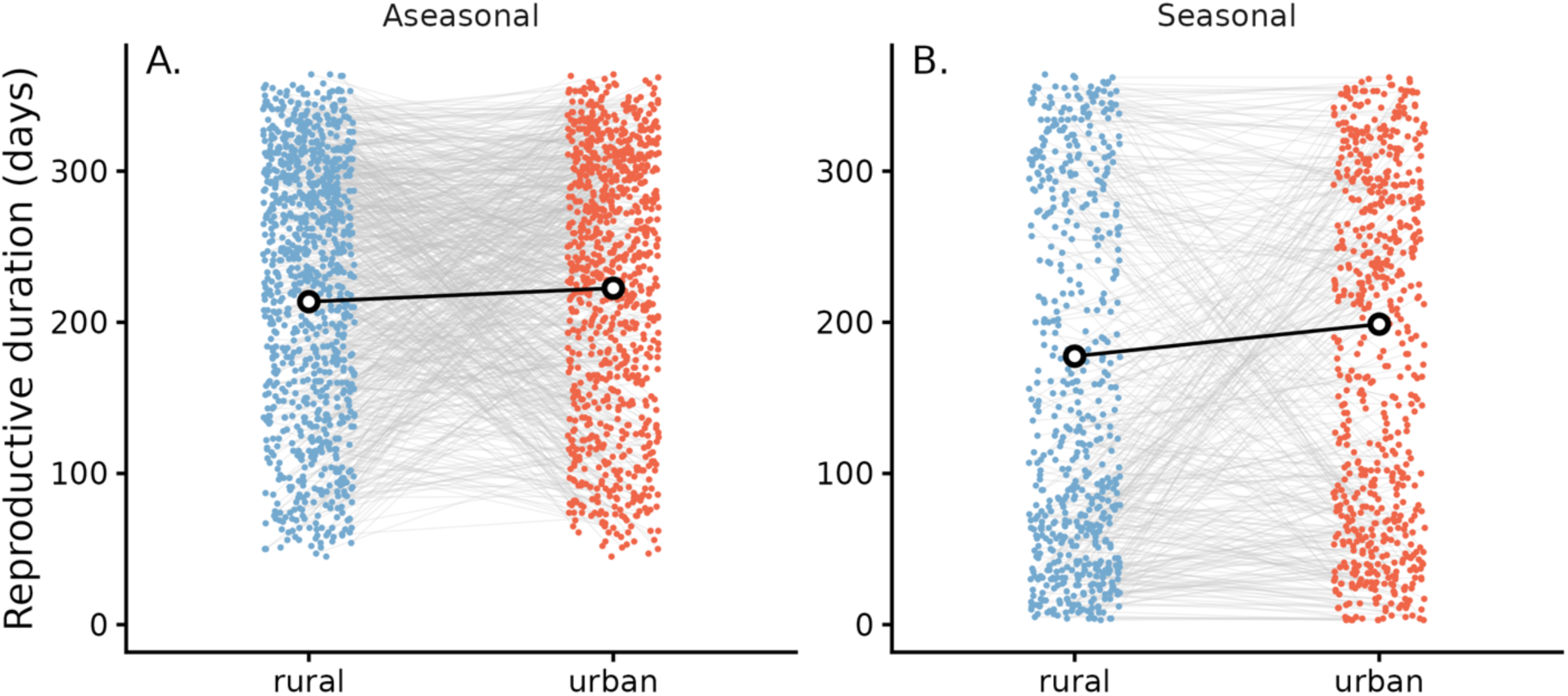
Paired urban–rural reproductive durations for aseasonal and seasonal species found in both rural and urban grid cells. A: Species categorized as aseasonal (with no peak) showed a longer flowering duration in urban settings. B: Species with seasonality (single-peak) in flowering also displayed a similar pattern with longer duration of flowering in urban regions. Thin gray lines denote raw paired data; black points and lines represent model-estimated marginal means.

#### 3.3.3. Comparison across annual and perennial species

For annual species, reproductive periods were nearly a month longer in urban than in rural localities (rural reproductive duration = 171.29 days ± 8.4; urban reproductive duration = 201.12 ± 8.15; p = 1.4 × 10^ (-8); Fig. 6A and Table S1). In contrast, reproductive periods were about a week longer in urban than in rural settings for perennial species (rural reproductive duration = 205.33 days ± 5.5; urban reproductive duration = 213.5 ± 5.46; p = 1.4 × 10^-8; Fig. 6A and Table S1). Similarly, annual plant species had considerably lower seasonality (*r*) in urban than in rural settings, but urbanization was associated with only a modest reduction in seasonality for perennial species (Fig. S3 and Table S2).

**Figure 6.**
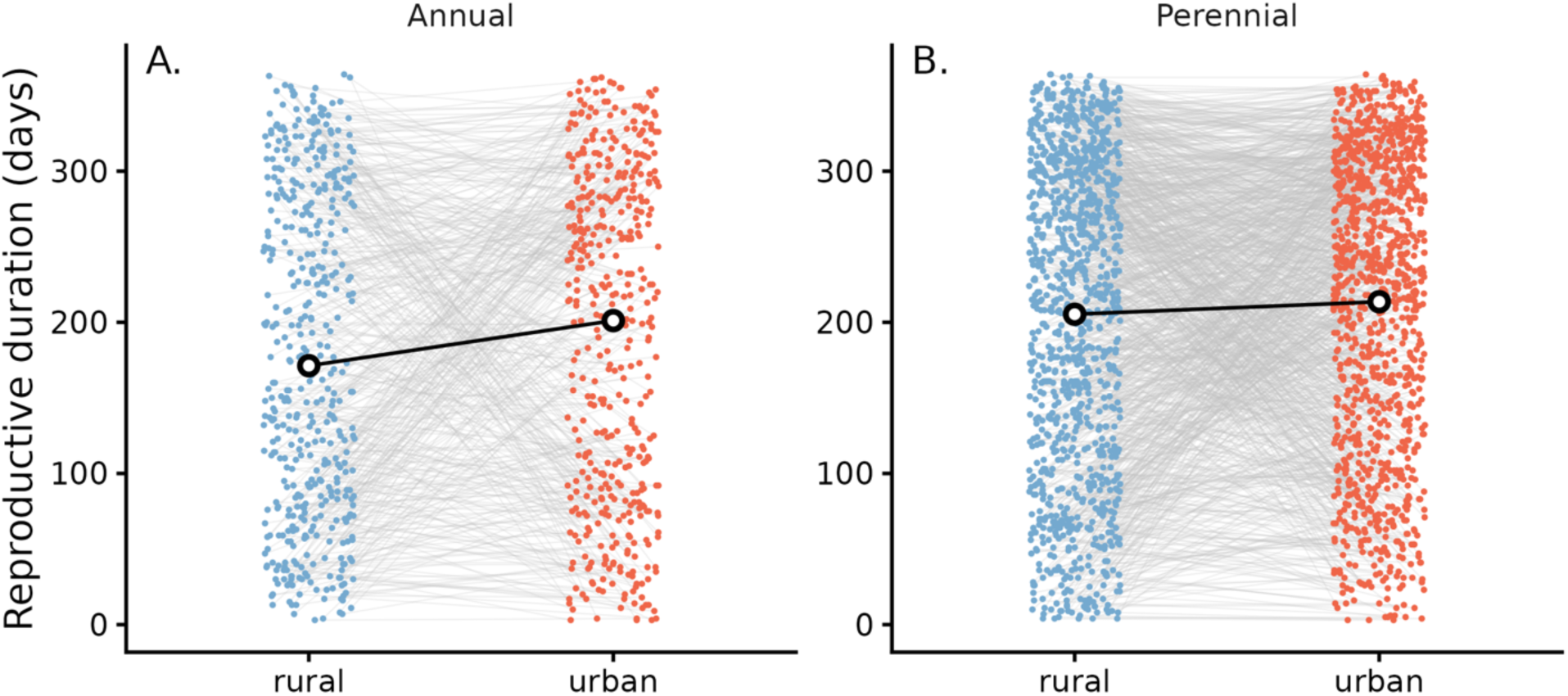
Paired urban–rural durations for annual and perennial species found in both rural and urban grid cells. A: Annual species displaying a significant longer reproductive phase in urban regions. B: Perennial species showing similar length of reproductive phase for rural and urban observations. Thin gray lines denote raw paired data; black points and lines represent model-estimated marginal means.

#### 3.3.4. Comparison across herbs, shrubs, and vines

Reproductive periods were significantly longer in urban than in rural localities for herbaceous plants and for vines (Herbaceous: rural reproductive period = 187.51 ± 6.61, urban reproductive period = 211.9 ± 6.47, p = 5.07e-10; Vines: rural reproductive period = 196.79±10.52, urban reproductive period = 212.25 ± 10.23, p = 3.49e-4, see Figs. 7A and 7C). However, no such effect of urbanization on reproductive period was found in shrubs (rural reproductive period = 209.43±6.89, urban reproductive period = 207.85 ± 6.76, Fig. 7B, n.s.)). Similar patterns were detected with the strength of seasonality r, which was lower in urban than in rural localities for herbs and vines but not for shrubs (Fig. S4 and Table S2).

**Figure 7.**
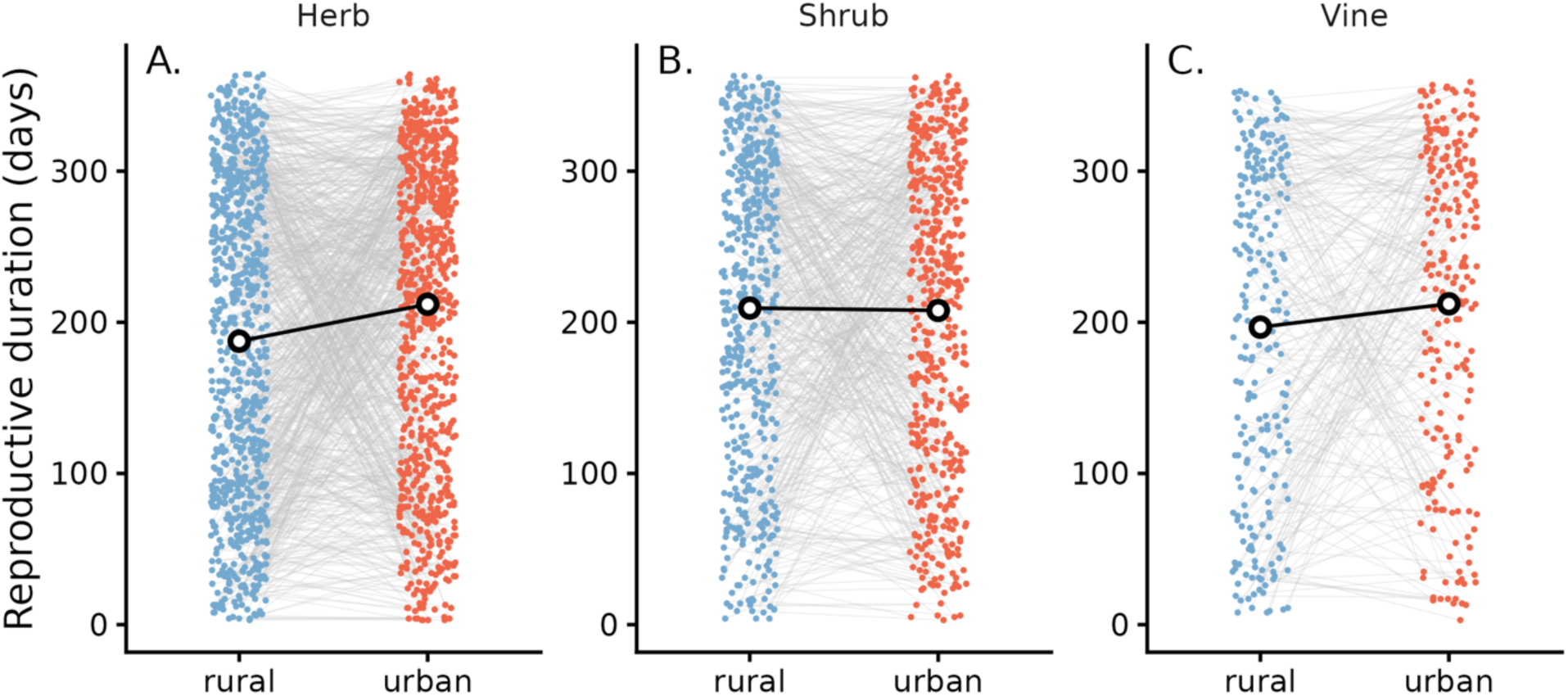
Paired urban–rural reproductive durations for herbs, shrubs, and vines found in both rural and urban grid cells. A: Herbs displaying a significant longer reproductive phase in urban regions. B: Shrubs showing no difference in reproductive duration between rural and urban observations. C: Vines with a significant difference in reproductive duration between rural and urban observations. Thin gray lines denote raw paired data; black points and lines represent model-estimated marginal means.

## 4. DISCUSSION

Urbanization is among the most important drivers of environmental change across the tropics, yet its influence on the timing of plant reproductive phenology remains poorly studied at broad spatial and taxonomic scales. By integrating multi-continental citizen science observations with curated herbarium records, our study reveals two central findings. First, despite fundamental differences in sampling design, citizen science and herbarium datasets converge on consistent estimates of species-level reproductive seasonality strength at local grid scales, supporting the use of citizen science-based observations for phenology-related studies in the tropics. Second, across thousands of species-grid combinations, urban plant populations exhibit longer reproductive duration and reduced seasonality strength relative to rural populations, although the magnitude and direction of this effect vary across species and functional groups. Together, these results demonstrate that urbanization leaves a detectable and biologically meaningful imprint on tropical reproductive phenology, while also highlighting the value of combining opportunistic and curated data streams to understand phenological responses to global change in the tropics.

### 4.1. Using citizen science data for studying tropical urban phenology

A vital contribution of our study lies in demonstrating the efficiency of citizen science observations in answering phenological questions that would be difficult to address using herbarium data alone. Although herbarium collections are taxonomically rich and provide vital phenological insights (Karthikeyan et al., 2025; Ordoñez et al., 2025), these observations tend to be from less developed (or rural) locations (Fig. 2b), which limits their utility for studying urban ecology. In contrast, iNaturalist observations are distributed over dense urban centers, peri-urban mosaics, and rural inland habitats, facilitating urban phenology studies. As a result, the use of iNaturalist data in peer-reviewed studies has increased rapidly, with the data coming from 128 countries representing 638 taxonomic families (Mason et al., 2025). This extensive coverage by iNaturalist illustrates the expanding reach of citizen science, even in biodiversity-rich but traditionally under-sampled tropical regions (Barve et al., 2020; Iwanycki Ahlstrand et al., 2022).

At the same time, the dataset from herbaria, though biased toward rural observations, offer taxonomically rich, high-quality records and remain equally important in phenology and climate change studies. This result (3835 unique species from herbarium records compared to 296 from iNaturalist) aligns with previous demonstrations that herbarium records capture high taxonomic and functional diversity (Eckert et al., 2024), which could make them reliable data sources for studying phenological signatures of diverse plant taxa (Iwanycki Ahlstrand et al., 2022; Willis et al., 2017). Our comparison of reproductive seasonality between data sources further reveals that many species classified as moderately and strongly seasonal based on herbaria records are referred as weakly seasonal in iNaturalist observations. This discrepancy in seasonality classification exists likely due to differences in sampling strategies. Herbarium specimens are often collected during peak phenophases via directed short field campaigns, whereas citizen science platforms like iNaturalist observations are results of opportunistic, repeated encounters across seasons. These contrasts between the herbaria and citizen science datasets affirm their complimentary strength, as combining these datasets bridges spatial and ecological gaps in tropical monitoring by integrating real-time, human-centered observations with long-term scientific records (Ramirez-Parada et al., 2024; Williamson et al., 2025). It also supports previous claims regarding the potential value of dedicated citizen science efforts in tropical areas, and particularly in tropical cities, to fill crucial data gaps in support of long term research and monitoring (e.g., SeasonWatch, Ramaswami et al., 2021).

### 4.2. Urbanization impacts on plant reproductive phenology

Analyzing nearly 3100 combinations of grid cells, species, and urban classes, we observed that urban populations had longer reproductive duration and lower seasonality strength than rural observations. Our observed patterns across the tropics mirror findings from temperate cities, where urban heat islands, increased atmospheric CO2, regulated irrigation, and altered light regimes prolong resource availability. These signatures of urbanization enable plants to flower earlier, flower longer, or produce multiple flowering cycles within a year (Fujiwara et al., 2025; Neil & Wu, 2006; D. S. Park et al., 2023; Sexton et al., 2023; Wohlfahrt et al., 2019). The similarity in phenological signal between temperate and tropic responses to urbanization may seem surprising, given the widely documented complexity and diversity of tropical phenology strategy. Here, the convergence between tropical and temperate urban phenology responses may reflect common urban mechanisms overriding otherwise distinct climate drivers. Indeed, our detection of a consistent signal of urbanization underscores its potentially powerful role as a driver of phenological shifts among diverse taxa.

Despite the clear urbanization effect in our global model (Table S1, S2), we also saw pronounced interspecific variation in both the direction and magnitude of phenological responses. Trait-level analyses (Figs.5-7) reveal varied phenological responses between life cycles and plant functional types. Seasonal and annual species, particularly with herbs and vines, exhibited stronger urban-associated increases in reproductive duration and reduction in seasonality strength. Longer reproductive periods for annual plants in urban settings might be due to their being able to exploit prolonged favorable conditions (Fujiwara et al., 2025), in contrast to perennial species, which are more likely to retain conservative phenological schedules (Marcacci et al., 2023; Sexton et al., 2023; Stanley & Ashman, 2025). Although relatively few studies have examined variation in phenological responses across functional groups (herbs, shrubs, vines) to urbanization, recent results from a review paper suggest that the composition of plant species and their functional group can lead to diverse phenology due to differences in growth characteristics and adaptability (Sun et al., 2026). As temperature is found to be a primary driver of flowering (Sun et al., 2026; Zhou et al., 2023), and temperature tends to vary between urban-rural settings (Ogunbode et al., 2025; Ribeiro et al., 2024), these variation in temperature and distinctiveness of plant species across functional group could explain variation in phenological responses across functional groups between urban-rural environments.

Together, the results suggest a clear effect of urbanization on tropical reproductive phenology through an extended reproducing duration in urban settings. Prolonged reproductive phases could increase overlap among co-flowering species, modify plant-pollinator interactions, alter mutualistic networks, and intensify interspecific pollen transfer, with consequences for reproductive success and competitive dynamics (Dzul-Cauich & Munguía-Rosas, 2025; Marcacci et al., 2023; Sexton et al., 2023). At the broader scale, lengthened reproductive seasons may have ecosystem-level consequences by altering carbon uptake, surface energy balance, and feedbacks between vegetation and urban microclimates (Williamson et al., 2025; Wohlfahrt et al., 2019). In rapidly urbanizing tropical regions, where biodiversity is high and ecological interactions are tightly linked to phenological timing, such shifts could propagate through mutualistic networks and ecosystem processes, meriting further study.

### 4.3. Limitations

Our work provides important insights into the implications of urbanization for plant phenology across the tropics, but a few important caveats should shape the interpretation of our results. Our analyses are implemented with urbanization classifications at a 1x1 km resolution within 25x25 km grid cells, assuming limited impacts of fine-scale environmental heterogeneity on the phenology within urban-rural landscapes. Our urban classification likewise simplifies a multidimensional gradient that includes variation in land use, management intensity, and microclimate. On-the-ground studies of phenology and demography across urban-rural gradients can help disentangle these multidimensional signatures of urbanization. Additionally, both citizen-science and herbarium datasets remain influenced by observer behavior and collection practices, particularly in urban settings where observations are more frequent and temporally biased. Although we control for potential biases towards higher density of sampling in urban settings (Fig. S5), it remains possible that more extensive temporal coverage of data contribute to a broader estimate of flowering duration or weaker seasonality strength in urban environments. Finally, our comparison of phenology estimates from iNaturalist, and herbarium specimens assumed that all herbaria specimens were reproductive at the time of collection. Although this assumption is consistent with most collection practices (Goëau et al., 2020; Heberling & Isaac, 2017), it may have led to our overestimating reproductive periods. Making herbarium specimens more valuable for phenological research will require continued advancements that draw on human and machine-based annotations of herbaria observations (Grady et al., 2025).

### 4.4. Conclusions

This study represents one of the first large-scale applications of iNaturalist to investigate plant phenology across urban contexts in tropical ecosystems. By coupling these records with herbarium observations, we demonstrate that opportunistic, crowd-sourced imagery can yield ecologically consistent and complementary insights into reproductive periodicity. The approach establishes a scalable framework for monitoring phenological resilience to urbanization and climate change, particularly in data-poor tropical regions where systematic observations are limited. Our findings suggest that abiotic features associated with urbanization result in extended reproductive periods than in nearby rural localities, with potential downstream impacts on population, community, and ecosystem dynamics. As digital archives and machine-learning-assisted phenophase annotation advance, integrating citizen-science, herbarium, and remote-sensing data will enable dynamic, high-resolution phenological mapping across space and time. Such efforts can transform tropical ecology from a historically data-scarce discipline into one driven by continuous, community-powered observation networks, improving forecasts of biodiversity responses to global change.

## 5. Author Contributions

Rohit Raj Jha, Anita Simha, Richard Ekeng Ita, Rachana Rao, and Gaurav Kandlikar conceived the idea. All authors contributed to data acquisition. Data curation was done by Rohit Raj Jha and Gaurav Kandlikar. Rohit Raj Jha completed the formal analysis with Gaurav Kandlikar. Rohit Raj Jha wrote the paper with inputs from Gaurav Kandlikar, Daijiang Li, and Anita Simha, and all authors reviewed the paper.

## Acknowledgements

We thank the contributors to iNaturalist and Herbarium observations and the authors of various open-source packages that make this computational work possible. We acknowledge the Louisiana State University startup funds to Gaurav Kandlikar and the National Science Foundation grant DBI2223508 to Daijiang Li. We thank Karthik Thrikkaderri for his insights during data acquisitions and comments on the manuscript.

## Ethics statement

None

## Conflicts of Interest

The authors declare no conflicts of interest.

## Data Availability Statement

The data and codes associated with this manuscript are archived in Zenodo (https://doi.org/10.5281/zenodo.18252047).

## Notes

https://doi.org/10.5281/zenodo.18252047

### Competing Interest Statement

The authors have declared no competing interest.

